# Direct-from-specimen microbial growth inhibition spectrums under antibiotic exposure and comparison to conventional antimicrobial susceptibility testing

**DOI:** 10.1101/2021.02.12.430910

**Authors:** Jade Chen, Su Su Soe San, Amelia Kung, Michael Tomasek, Dakai Liu, William Rodgers, Vincent Gau

**Author notes:** corresponding author, (VG).

## Abstract

Increasing global travel and changes in the environment may increase the frequency of contact with a natural host carrying an infection, and therefore increase our chances of encountering microorganisms previously unknown to humans. During an emergency (man-made, natural disaster, or pandemic), the etiology of infection might be unknown at the time of patient treatment. The existing local or global Antimicrobial Stewardship Programs might not be fully prepared for emerging/re-emerging infectious disease outbreaks, especially if they are caused by an unknown organism, engineered bioterrorist attack, or rapidly evolving superbug. We demonstrate an antimicrobial efficacy profiling method that can be performed in hours directly from clinical urine specimens. The antimicrobial potency is determined by the microbial growth inhibition and compared to conventional antimicrobial susceptibility testing (AST) results. The oligonucleotide probe pairs on the sensor were designed to target gram-negative bacteria, specifically *Enterobacterales* and *Pseudomonas aeruginosa*. A total of 10 remnant clinical specimens from the CLIA labs of New York-Presbyterian Queens were tested, resulting in 100% categorical agreement with reference AST methods (Vitek and broth microdilution method). The combined categorical susceptibility reporting of 12 contrived urine specimens was 100% for ciprofloxacin, gentamicin, and meropenem over a range of microbial loads from 10^5^ to 10^8^ CFU/mL.

## Introduction

Direct-from-specimen microbial growth inhibition assessment can assist in emergency preparedness and pre-hospital interventions with timely patient-specific antimicrobial efficacy profiling information. The very concept of empirical therapy is a testament to the reality that the current methods used in clinical microbiology labs are often unable to render information in a time frame that can inform initial treatment decisions [1]. Phenotypic antimicrobial efficacy profiling, where clinical specimens are directly exposed to different antibiotic conditions, could provide critical information for the prescription of antibiotics in hours. The results of a phenotypic antimicrobial efficacy profile test, taken in conjunction with local antibiogram data, could guide the course of therapy to improve patient outcomes and slow the spread of antimicrobial resistance. We demonstrate a molecular test based on the transcriptional responses of causative bacteria to antibiotic exposure directly from urine specimens. Quantification of group-specific or species-specific 16S rRNA growth sequences is used to provide rapid antimicrobial efficacy profiling results, bypassing the necessity of overnight culture for generating isolates. Categorical agreement is assessed with reference AST methods according to CLSI guidelines.

Even though antibiotics do not directly affect the SARS-CoV-2 respiratory virus responsible for the COVID-19 pandemic, physicians are administering many more antibiotics than normal when treating COVID-19 patients [2]. As published in the New England Journal of Medicine, a majority of the surveyed 1,099 COVID-19 patients (58.0%) received intravenous antibiotic therapy in China, while only 35.8% received oseltamivir antiviral therapy [3]. Antibiotic use appears to be surging and higher percentages of COVID-19 patients with severe conditions and pediatric patients (88% in a multicenter pediatric COVID-19 study [4]) received antibiotic therapies. WHO warned that the majority of COVID-19 patients in the U.S. and Europe received similar antibiotic treatments from physicians.^5^ Because viral respiratory infections often lead to bacterial pneumonia, physicians can struggle to identify which pathogen is causing a person’s lung problems. A recent study by Zhou et al. [6] found that 15% of 191 hospitalized COVID-19 patients - and half of those who died - acquired bacterial infections. Major outbreaks of other respiratory viruses illustrate the same concern: the majority of deaths from the 1918 flu showed autopsy results consistent with bacterial pneumonia, and up to half of the 300,000 people who died of the 2009 H1N1 flu were confirmed to have died from pneumonia [7–8]. Therefore, a shorter time to rule out certain antibiotic options if there is microbial growth under such conditions can provide the physicians valuable information before the availability of conventional AST results.

Microbial growth inhibition response curves to antibiotic exposure conditions across a range of microbial loads can provide a dynamic and rapid method for estimating antimicrobial efficacy in a much shorter timeframe than the endpoint minimum inhibitory concentration (MIC) method used in conventional AST. Here, we present a method to quantify the 16S rRNA content of viable targets pathogens in raw specimens such as urine following exposure to certain concentrations of an antibiotic *in vitro*, and we have developed a method to interpret the antimicrobial effect by analyzing the differential microbial responses at two dilutions. The hypothesis is that the growth inhibition concentration (GIC) is the lowest concentration necessary to inhibit growth in all strains in a given sample after adjusting for pathogen concentration effects. We compare the GIC reported from this antimicrobial efficacy profiling directly from the specimen with the MIC and susceptibility reporting from CLSI reference methods to obtain the categorical agreement, and we then establish a correlation between the microbiological susceptibility (i.e., MIC) and antimicrobial efficacy (i.e., GIC).

### Electrochemical-based molecular quantification of RNA transcription for streamlined ID and phenotypic AST

Prior to developing antimicrobial efficacy profiling directly from unprocessed specimens, a PCR-less RNA quantification protocol through enzymatic signal amplification with a proprietary electrochemical sensor array was developed, applied to streamlined pathogen identification and AST with species-specific probe pairs, validated and published with contrived and remnant clinical specimens with our clinical collaborators [9–44]. The detection strategy of our universal, electrochemical-based sensors is based on sandwich hybridization of capture and detector oligonucleotide probes which target 16S rRNA. The capture probe is anchored to the gold sensor surface, while the detector probe is linked to horseradish peroxidase (HRP). When a substrate such as 3,3’,5,5’-tetramethylbenzidine (TMB) is added to an electrode with capture-target-detector complexes bound to its surface, the substrate is oxidized by HRP and reduced by the bias potential applied onto the working electrode. This redox cycle results in shuttling of electrons by the substrate from the electrode to the HRP, producing enzymatic signal amplification of current flow in the electrode. The concentration of the RNA target captured on the sensor surface can be quantified by the reduction current measured through the redox reaction between the TMB and HRP with a built-in multi-channel potentiostat in our system. The implementation of robotic automation of the molecular quantification of 16S rRNA transcription as a growth marker on the current lab automation system was to address the adaptation into the workflow of a clinical microbiology laboratory and the assay variance caused by manual operation [45]. The centrifugation-based specimen preparation can be performed on our current systems, but manual specimen processing was used in this study for assay parameter optimization. The change in RNA transcription is among the earliest cellular changes upon exposure to antibiotics, long before phenotypic changes in growth can be observed [46]. Quantifying changes in RNA signatures is therefore a particularly appealing approach for slow-growing organisms [47]. Measuring the RNA response of pathogens to antibiotic exposure directly in clinical specimens would provide a rapid susceptibility assessment that can be performed in clinical settings.

## Material and Methods

### Bacterial strains and antibiotic stripwells

Strains included in this study were obtained from various sources including the CDC AR Bank and New York-Presbyterian Queens (NYPQ) and consisted of the following organisms listed with the number of clinical isolates: 11 *Escherichia coli*, 5 *Klebsiella pneumoniae*, and 5 other species as detailed in S1 Table. All clinical isolates were obtained anonymously from remnant patient samples collected for routine culture and were de-identified prior to testing under the approved NYP/Queens Institutional Review Board and joint master agreement. We aimed to test an even distribution of species with MIC values on or near the susceptible and resistant breakpoints of each antibiotic including three representative antibiotics of three different classes (fluoroquinolones, aminoglycosides, and carbapenems): ciprofloxacin (CIP; Cayman Chemical Company, Ann Arbor, MI), gentamicin (GEN; Sigma-Aldrich, St. Louis, MO), and meropenem (MEM; Cayman Chemical Company). CDC AR Bank isolates were used to include representative bacteria susceptibility profiles that were not covered by those from NYPQ. CDC AR Bank isolates were stored as glycerol stocks at −80°C and were grown from these stocks at 35°C on tryptic soy agar plates with 5% sheep’s blood (Hardy Diagnostics) for 18-24 hours before testing. Suspensions of each isolate to be used for contriving urine samples were prepared using cation-adjusted Mueller-Hinton II broth and a Grant DEN-1B densitometer (Grant Instruments, Cambridge, UK). Negative urine specimens to be used for testing of contrived samples were stored in Falcon tubes at 4°C. Clinical urine samples from NYPQ were stored in BD 364954 Vacutainer Plus C&S tubes containing boric acid at 4°C prior to overnight shipment for testing. Consumables consisted of stripwells with dried antibiotics, electrochemical-based sensor chips functionalized with oligonucleotide probe pairs complementary to *Enterobacterales* and *Pseudomonas aeruginosa* for RNA quantification, and a reagent kit for lysing and viability culture. Stripwells were prepared by drying antibiotics in DI water with 0.1% Tween onto EIA/RIA 8-well strips (Corning, Corning, NY) at the following concentrations: CIP 0.0625, 0.125, 0.25, 0.5, 1, 2, 4 μg/mL; GEN 1, 2, 4, 8, 16, 32 μg/mL; MEM 0.5, 1, 2, 4, 8, 16, 32 μg/mL. The first well of each stripwell was left without antibiotic to be used as a growth control (GC) during the assay.

### Specimen collection and matrix removal

Urine samples were spun down to remove the majority of matrix components in the supernatant. Specifically, urine samples with 4-mL starting volume were spun in a centrifuge at 5,000 RPM for 5 minutes, after which supernatant was removed and replaced with cation-adjusted MH broth to make 1x and 0.1x inoculums for delivery to the antibiotic exposure stripwells.

### Electrochemical-based microbial growth quantification

Since there are no commercially available FDA-cleared systems or CLSI reference methods to provide AST results directly from specimens without overnight culture or clinical isolates, the direct-from-specimen antimicrobial efficacy profiling approach presented in this study aims to demonstrate a significant correlation to conventional AST results. The electrochemical-based biosensor measures the reduction current from cyclic enzymatic amplification of an HRP label with TMB and H2O2. The resulting reduction current signal can be estimated with the Cottrell equation [48]. Signal levels (in nanoamps) from each microbial exposure well (no antimicrobial for GC well) were normalized to the one from the GC well and plotted against the spectrum of antimicrobial tested. Two antibiotic exposure stripwells with a spectrum of seven antibiotic concentrations and one GC were used for each specimen at 1x (undiluted pellet) and 0.1x (diluted pellet) to generate two microbial responsive curves. Each dual-response-curve signature was generated by overlaying two GC ratio curves over the antibiotic spectrum, establishing a signature library corresponding to each antimicrobial efficacy and microbial susceptibility combination. Changes in response signature and inflection point in GC curve were analyzed by three algorithms as in the corresponding tables in Supplemental Material to match a categorical classification (susceptible, intermediate, or resistant).

### Antibiotic exposure stripwell inoculation and molecular quantification

One hundred microliters of reconstituted specimen pellets (1x and 0.1x) were inoculated into each well of an AST stripwell. All stripwells were incubated at 35°C for the exposure time indicated in each study. Thirty-six microliters of 1M NaOH were added to each well to lyse target gram-negative pathogens after antibiotic exposure with a 3-minute incubation at room temperature. Twenty-four microliters of 1M HCl were then added to each well to neutralize the pH of the lysed sample, or lysate, and prevent the degradation of free RNA. Ten microliters of the lysate from each well were pipetted to its corresponding sensors on two electrochemical sensor chips (a total of 4 sensors per well). No sample was delivered to the negative control sensors. All chips were incubated for 30 minutes at 43°C, and the RNA content was quantified for microbial growth response as described above.

### Clinical feasibility validation with blind clinical specimens

The clinical specimens for the blind testing study were remnant specimens collected at NYPQ under the current IRB. Incoming urine specimens for urine culture as part of routine care with confirmed positives for either *Enterobacterales* or *Pseudomonas aeruginosa* were shipped overnight to GeneFluidics for testing. De-identification and data analysis were performed by administrative staff. Species belonging to the *Enterobacterales* family such as *Escherichia coli, Klebsiella* spp., and *Enterobacter* spp. are the major cause of urinary tract infections, blood-stream infections, and healthcare-associated pneumonia [49–50]. The *Enterobacterales* family and *Pseudomonas aeruginosa* were selected due to their increasing resistance to commonly used antimicrobial agents [51].

### Statistical analysis

Signals generated from each sensor from enzymatic reaction with TMB substrate were analyzed with three different algorithms for comparison. Before reporting GC ratio, the algorithm first assessed the signal level from the negative and growth controls from each sensor chip. If either control was out of the acceptable range (i.e., greater than 50 nA for the negative control, less than 50 nA for the growth control), the algorithm reported “NC fail” or “GC fail”, respectively, indicating substandard quality of a sensor chip or no bacterial growth. If all controls passed the acceptance criteria, the algorithm proceeded to determine the inflection point from the GC ratio plot against the antibiotic spectrum. The antibiotic concentration corresponding to the inflection point was estimated by two algorithms (inhibited growth cutoff and maximum inhibition) and reported as the growth inhibition concentration (GIC). The inhibited growth cutoff method reported the highest antibiotic concentration with a GC ratio lower than a predetermined cutoff value, so the GIC was determined solely based on the signal level from each antibiotic exposure condition normalized to the one from the growth control. Initial assessment used both 0.4 and 0.5 as cutoff values with on-scale strains to determine the final cutoff value. The maximum inhibition method reported the GIC as the highest antibiotic concentration after the maximum GC reduction in a microbiological response plot against a series of 2-fold dilutions of the antibiotic of interest, so the GIC corresponded to the greatest change in the slope of the response curve as a whole instead of signal levels. If the GC value from the lowest antibiotic concentration was less than 0.45, indicating significant growth inhibition, the GIC was reported as less than the lowest antibiotic concentration tested. If the GC value from the highest antibiotic concentration was higher than 0.9, indicating very limited growth inhibition, the GIC was reported as larger than the highest antibiotic concentration tested. The first level of analysis was qualitative, whereby the antimicrobial efficacy profiles (significant growth, moderate growth, and inhibited growth) derived from the GIC were compared to the corresponding antibiotic susceptibility results (R for resistant, I for intermediate, or S for susceptible) from the clinical microbiology lab or CLSI reference methods.

Any direct-from-specimen antimicrobial efficacy profiles found to be misclassified (i.e., GIC higher than the susceptible breakpoint for a susceptible strain) were retested with both growth inhibition and microdilution reference methods. Categorical agreements were calculated for each specimen type. As a second level of analysis for GIC to MIC comparison only, the discrepant GIC/MIC values (i.e., GIC 2-fold above or below the MIC value from the clinical microbiology lab) were retested and compared to the microdilution reference method. Essential agreements were calculated for each specimen type.

## Results

There is always a valid concern about the detection sensitivity and matrix interference when developing a direct-from-specimen microbial growth inhibition test without the need of an overnight cultured isolate. Since the direct-from-specimen microbial growth inhibition test starts with a specimen with unknown pathogen concentration from 0 to > 10^8^ CFU/mL in different specimen types, the correlation between the limit of detection (LOD) of the current molecular analysis platform with the assay turnaround time (TAT) was established in Fig 1 in order to determine the minimum assay time needed for quantification of RNA transcription at different levels of pathogen concentrations. As shown in Fig 1C, the TAT and dynamic range of ID can be configured to be from 16 minutes to 36 minutes by adjusting the analyte incubation time for higher target LODs. Target pathogen enrichment and matrix component removal can be carried out by centrifugation to achieve lower target LODs with TAT of 42 minutes to 110 minutes. For low-abundance pathogens and early infection diagnostics, additional viability culture steps with TAT of 4 to 5.5 hours can be included to achieve an LOD of < 10 CFU/mL. The direct-from-specimen antimicrobial efficacy profiling protocol was based on these assay parameters summarized in Fig 1D.

**Fig 1.**
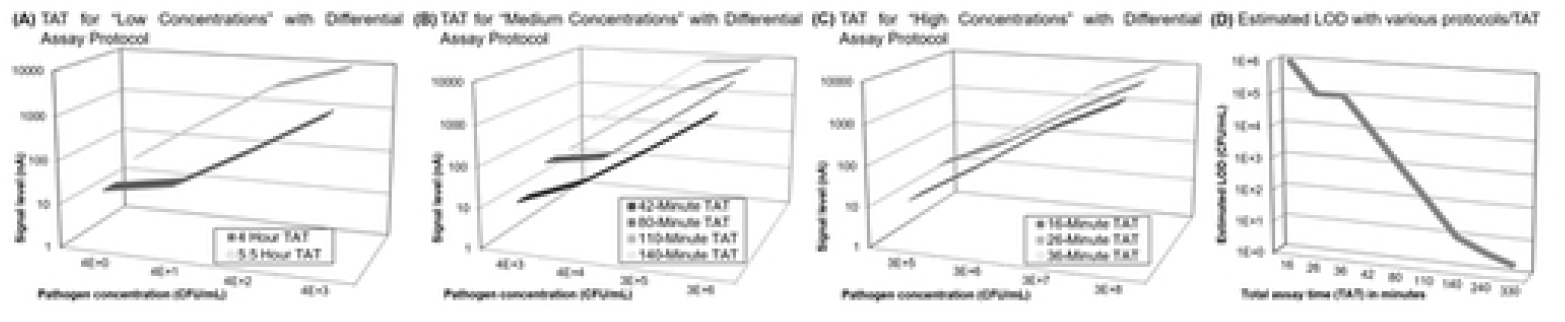
Calibration curves of configurable ID protocols with various TAT and LODs. (A) TAT for “low” pathogen concentrations, (B) “medium” pathogen concentrations, (C) “high” pathogen concentrations. (D) Summary of various TAT and LOD.

We first focus on the feasibility of assessing microbial growth inhibition without the potential interference of matrix effects by using contrived samples in culture media (Mueller Hinton Broth, Sigma-Aldrich) with one of two clinical isolates with distinct susceptibilities. The initial evaluation was conducted with highly susceptible *E. coli* (EC69, MIC ≤ 0.06 μg/mL for ciprofloxacin) and highly resistant *K. pneumoniae* (KP79, MIC: >8 μg/mL for ciprofloxacin) strains from the CDC AR Bank (Fig 2). Since the goal of the pilot study was to investigate the potential interference in urine, the spiked concentration was set to 10^7^ CFU/mL. Three antibiotic exposure times (30, 60, and 90 minutes) were tested as primary parameters for optimization. Microbial growth inhibition was plotted with the signal ratios normalized to the one from the growth control, GC ratio, against the ciprofloxacin concentrations tested ranging from 0.0625 μg/mL (two 2-fold dilutions under the *Enterobacterales* susceptible breakpoint) to 4 μg/mL (two 2-fold dilutions above the *Enterobacterales* resistant breakpoint). As shown in Fig 2A, all microbial response curves of resistant *K. pneumoniae* CDC 79 (no-fill pattern) were overlapping at the GC ratios at around 1.0 (see S2 Table for GIC reporting from all three algorithms), indicating no inhibited growth no matter how long the exposure time was. However, there was a clear trend of inhibited growth with lower GC ratios with the susceptible *E. coli* CDC 69 (gradient pattern), indicating more significant inhibited growth while increasing the exposure time or ciprofloxacin concentration. The reported GIC value from the Maximum Inhibition algorithm (see Algorithm section for details) is listed to the right of each response curve. We then repeated the same protocols with the contrived urine samples to evaluate the impact of the urine matrix components in Fig 2B. The bolded GIC value (S strain in MH 30 min, S strain in urine 30 min, S strain in urine 60 min) represents incorrect categorical susceptibility reporting, which occurs when the exposure time is insufficient. The microbial growth inhibition curves from contrived urine samples in Fig 2B exhibit identical characteristics as those in culture media in Fig 2A. This suggests the supernatant removal step is efficient enough to remove urine matrix, but not too harsh to put the pathogen into the stationary phase.

**Fig 2.**
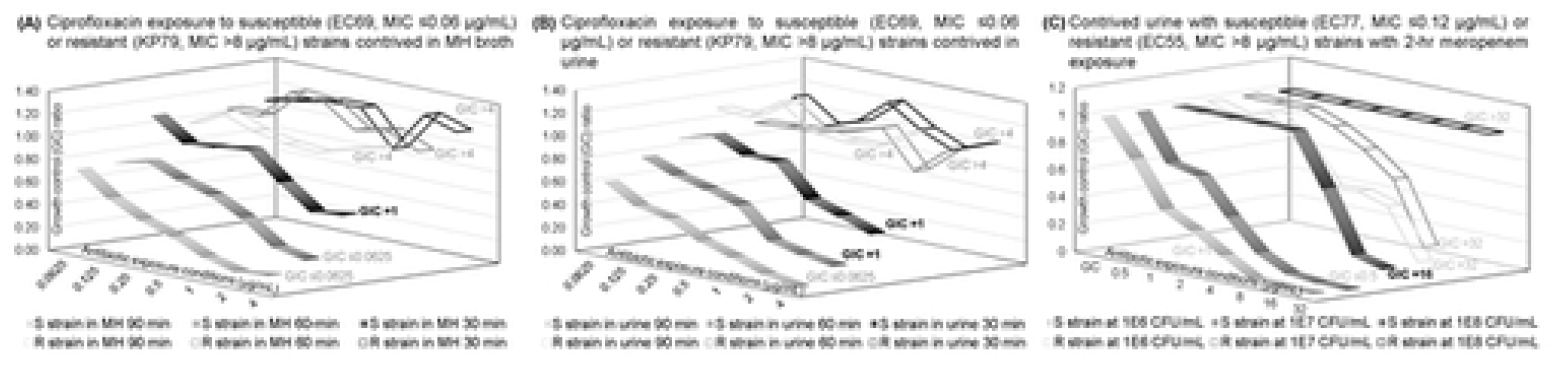
Investigation of matrix interference components and starting inoculum concentration. (A) Ciprofloxacin antimicrobial efficacy profiling in MH broth and (B) direct-from-urine ciprofloxacin antimicrobial efficacy profiling using very susceptible (*E. coli* CDC 69) and very resistant (*K. pneumoniae* CDC 79) strains from the CDC AR bank. (C) Direct-from-specimen meropenem antimicrobial efficacy profiling with 2-hr exposure for urine with very susceptible (*E. coli* CDC 77) and resistant strains (*E. coli* CDC 55). Bolded GIC values indicate incorrect categorical susceptibility as in short exposure time (30 or 60 min.) in Figs 2A and 2B or high microbial load (10^8^ CFU/mL) in Fig 2C.

Since a shorter antimicrobial exposure time might result in errors in categorical susceptibility reporting due to insignificant growth inhibition of susceptible strains compared to resistant ones as shown in Figs 2A and 2B without a more sophisticated algorithm, we suspected the similar insignificant growth inhibition separation could occur if the microbial load is much higher than the standard inoculum density of 5×10^5^ CFU/mL. We needed to adjust the antibiotic exposure time and the matrix interference reduction procedures for each specimen types for the maximum differential antimicrobial efficacy profiling between susceptible and resistant strains over a physiological range of microbial loads. Contrived urine samples were used at three different microbial loads against a different class of antibiotic. To explore biological, chemical and molecular analytical limitations, shorter antibiotic exposure times were used for urine samples in Fig 2C to assess the separation of responses curves from both resistant and susceptible strains. Antimicrobial efficacy profiling tests directly from urine contrived samples were evaluated. Based on the trend of GC ratio changing along the increasing meropenem concentrations (0.5 to 32 μg/mL), the GIC would be reported as “susceptible” (≤ S-breakpoint of 1 μg/mL for meropenem) for *E. coli* CDC 77 (MIC: ≤ 0.12 μg/mL) and “resistant” (≥ R-breakpoint of 4 μg/mL for meropenem) for *E. coli* CDC 55 (MIC: > 8 μg/mL), which agree with the categorical susceptibility from CDC AR Bank even though the reported GIC was not exactly the same as the MIC value. To establish a higher correlation between the MIC and GIC values it would be necessary to incorporate the impact of inoculum effect on the GIC reporting, which is not within the scope of this initial study. With higher contrived concentrations, we expect the inflection point would be higher due to the higher bug-to-drug ratio. Even for susceptible strains, microbial growth can be observed at low antibiotic exposure concentrations at or below the susceptible breakpoint if the microbial load is higher than the typical inoculation concentration at 5×10^5^ CFU/mL.

Because Fig 2C only demonstrated the feasibility to differentiate highly susceptible from highly resistant strains, which do not represent all clinical strains, we wanted to evaluate the growth inhibition curves with on-scale strains (with an MIC value on or near the susceptible or resistant breakpoints) with *E. coli* CDC 1 with an MIC of 4 μg/mL for gentamicin (on susceptible breakpoint), *E. coli* CDC 85 with an MIC of 1 μg/mL for meropenem (on susceptible breakpoint), *K. pneumoniae* CDC 80 with an MIC of 0.5 μg/mL for ciprofloxacin (on intermediate breakpoint) and the exposure time at 2, 3 and 4 hours. General susceptibility trends with inhibited growth were observed at 2, 3 and 4 hours as shown in Fig 3 with different slopes along the antibiotic exposure conditions (1 to 32 μg/mL for gentamicin, 0.5 to 32 μg/mL for meropenem, and 0.0625 to 4 μg/mL for ciprofloxacin). The GIC values reported for *E. coli* CDC 1 were all at 2 μg/mL for all exposure times, and they were within one two-fold dilution of the MIC (see S3 Table for GIC reporting from all three algorithms). In addition, the categorical susceptibility indicated in the parentheses is correctly reported as susceptible. The GIC values for *E. coli* CDC 85 moved up from ≤0.5 to 2 μg/mL when meropenem exposure time was increased from 2 to 4 hours. Even though the GIC values at all three exposure times were within one dilution of the MIC for *E. coli* CDC 85, longer exposure times strongly align the GIC with the MIC value. The reproducibility of GIC reporting from two different batches of stripwell (010621 and 112420) was evaluated in Figs 3C and 3D, and the GIC reporting was consistent at all three conditions. The initial GIC reporting with just two hours of ciprofloxacin exposure was 0.5 μg/mL for both batches, and it agreed with the MIC values from CDC AR Bank database. However, the GIC value transitioned to 0.125 μg/mL with longer exposure time. The MIC from the microdilution method for *K. pneumoniae* CDC 80 was 0.25 μg/mL, which is within one two-fold dilution from the GIC reporting in all exposure times with both batches of stripwell.

**Fig 3.**
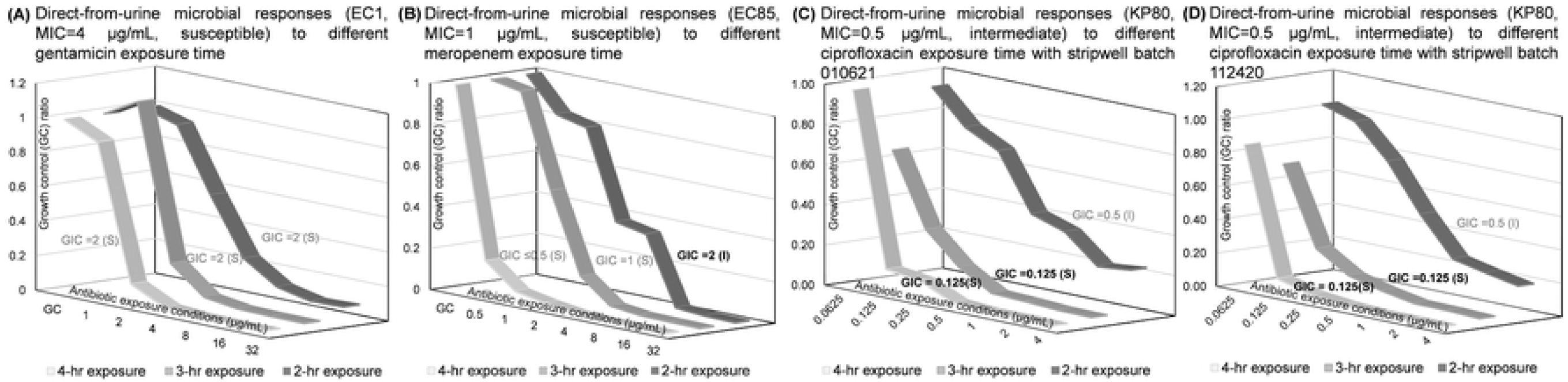
Varying antibiotic exposure times for direct-from-urine antimicrobial efficacy profiling of on-scale strains for different antibiotics classes. (A) Gentamicin responses. (B) Meropenem responses. (C, D) Ciprofloxacin responses with different stripwell batches. Bolded GIC values indicate incorrect categorical susceptibility.

After demonstrating that 3 hours of exposure time is sufficient for determining categorical susceptibility based on the reported GIC compared to the one based on the MIC reported from the microdilution reference method, we explored the ability to differentiate bacterial strains with on-scale MIC values over the range on or near the susceptible and resistant breakpoints in Fig 4. Fig 4A shows the growth inhibition responses to ciprofloxacin from *E. coli* (EC69: MIC≤0.0625 μg/mL, EC85: MIC> 8μg/mL) and *K. pneumoniae* (KP126: MIC= 0.125μg/mL, KP80: MIC= 0.5 μg/mL, KP76: MIC= 1μg/mL), and there is a clear trend in the shift of GIC (from ≤0.06 μg/mL to >4 μg/mL) along with the MIC values (from ≤0.06 μg/mL to >8 μg/mL). See S4 Table for GIC reporting from all three algorithms. Similarly, the growth inhibition responses to gentamicin from *K. pneumoniae* (KP126: MIC≤ 0.25μg/mL, KP79: MIC >16 μg/mL) and *E. coli* (EC1: MIC=4 μg/mL, EC451: MIC=8 μg/mL, EC543: MIC=16 μg/mL) are illustrated in Fig 4B, and there is a clear trend in the shift of GIC (from ≤1 μg/mL to >32 μg/mL) along with the MIC values (from ≤0.25 μg/mL to >16 μg/mL). The categorical susceptibility of all susceptible and resistant strains was reported correctly based on the reported GIC value except the two intermediate strains (KP80 for ciprofloxacin and EC451 for gentamicin). Both intermediate strains reported a GIC value two-fold lower than the MIC values from the reference methods. The essential agreement based on MIC/GIC values are acceptable, but both intermediate strains had minor errors according to the CLSI M100 and FDA Class II Special Controls Guidance for AST systems [52–53].

**Fig 4.**
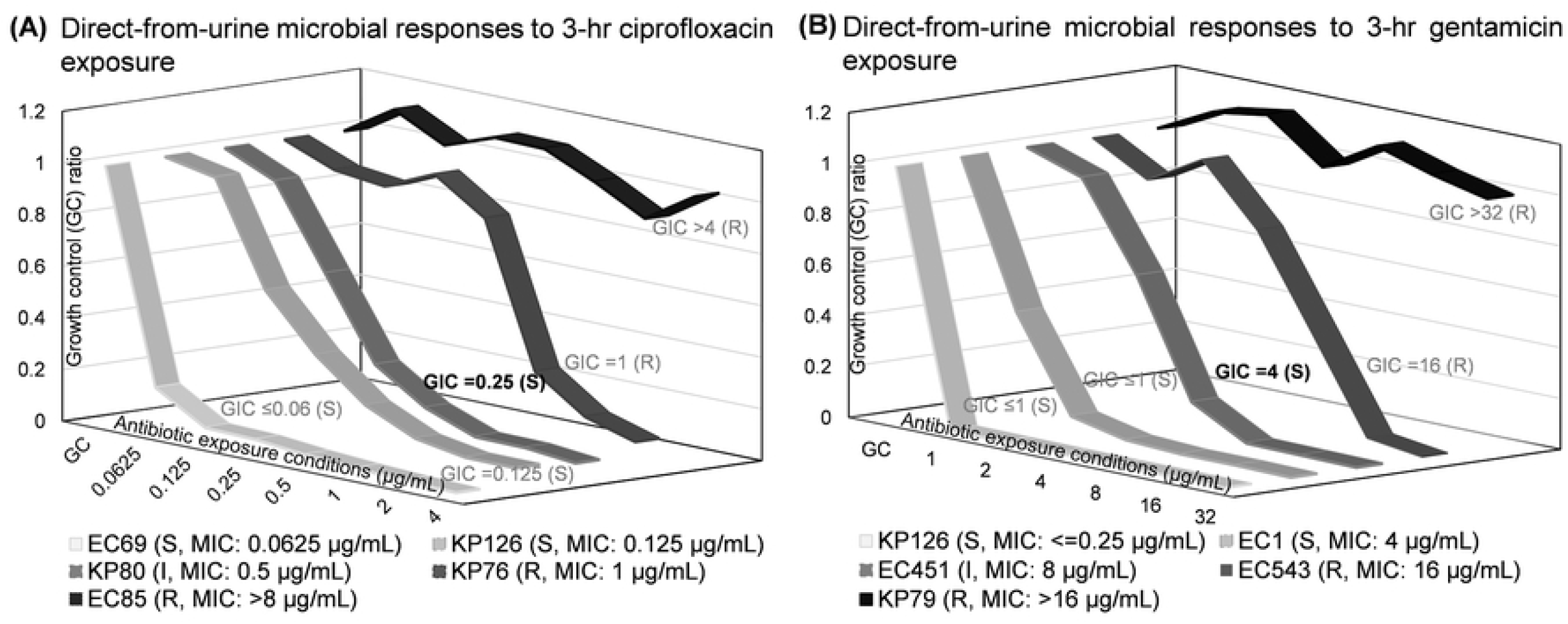
Direct-from-urine antimicrobial efficacy profiling of pathogens with a range of on-scale MIC values. All urine specimens contrived at 10^6^ CFU/mL. (A) Ciprofloxacin responses. (B) Gentamicin responses. Bolded GIC values indicate incorrect categorical susceptibility.

As shown in bolded GIC values, categorical susceptibility reporting (susceptible, intermediate or resistant) might be incorrect if the antimicrobial exposure time is too short (Figs 2A, 2B, and 3B), microbial load is too high (Fig 2C), or the MIC is on intermediate or resistant breakpoints (Figs 3C-D and 4A-B). Besides extending the antimicrobial exposure time, especially for time-dependent antibiotics such as meropenem, we explored the feasibility of a dual-kinetic response approach to cover a broader range of microbiological responses by inoculating two sets of seven antimicrobial concentrations in two antibiotic stripwells with clinical specimens at the original concentration (1x) and the 10-fold dilution (0.1x). Additionally, to ensure that the current GIC reporting algorithm is correlated with the microbial susceptibility and MIC values throughout the physiological range, a set of microbial loads in urine (10^5^ to 10^8^ CFU/mL) were tested in Fig 5, and the GIC was calculated from the dual kinetic (1x and 0.1x) curves. The inflection point shifted toward higher antibiotic concentrations with higher microbial loads, but the GICs were the same without the inoculum effect adjustment (see S5 Table for GIC reporting from all three algorithms). In Fig 5B, the growth inhibition curves of 1x and 0.1x of 10^6^ CFU/mL overlap with each other in the insert graph, even though the signal levels of these two sets of curves were very different. Because the microbial load from 10^5^ to 5×10^5^ CFU/mL and from 5×10^5^ to 10^6^ CFU/mL is the same (2-fold), the symmetrical characteristics result in overlapping GC ratio curves. Figs 5A-D show the transition of GIC reporting from ≤0.0625 μg/mL (susceptible), 0.125 μg/mL (susceptible), to 1 μg/mL (resistant). The categorical susceptibility reporting of “susceptible” was correct over a range from 10^4^ CFU/mL (0.1x of 10^5^ CFU/mL) to 10^7^ CFU/mL (0.1x of 10^8^ CFU/mL). The GIC value jumped from 0.125 μg/mL (0.1x of 10^8^ CFU/mL) to 1 μg/mL (1x of 10^8^ CFU/mL) as shown in Fig 5D. Similar microbial responses were observed in the rapid ciprofloxacin exposure study with the same *E. coli* CDC 69 strain in Fig 2B; the GIC value jumped from 0.0625 μg/mL (90-min exposure) to 1 μg/mL (30-min and 60-min exposure). The GC signal levels as shown in S5 Table were saturated at 10,000 nA for 10^7^ and 10^8^ CFU/mL, so the reported GIC value is expected to be higher than the MIC values due to the inoculum effect.

**Fig 5.**
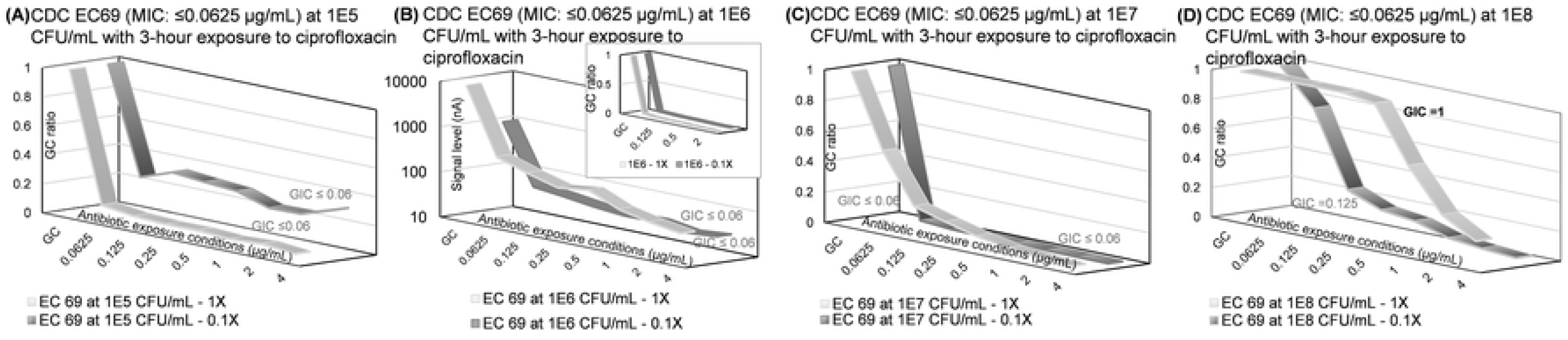
Direct-from-urine ciprofloxacin antimicrobial efficacy profiling with dual kinetic curves on different contrived urine concentrations. Dual kinetic curves for *E. coli* CDC 69 with MIC of ≤0.0625 μg/mL at starting sample concentrations of (A) 10^5^ CFU/mL, (B) 10^6^ CFU/mL, (C) 10^7^ CFU/mL, (D) 10^8^ CFU/mL. The bolded GIC value indicates incorrect categorical susceptibility.

The combined categorical susceptibility reporting as shown in Table 1 of the dual-kinetic-curve response in Fig 5 using the Maximum Inhibition algorithm is the maximum GC reduction in both microbiological response plots, so the combined GIC corresponds to the greatest change in the slope of both response curves. Table 1 is the summary of the individual and combined GIC reporting from all contrived concentrations in Fig 5. Since the combined categorical susceptibility is determined by the largest GC ratio change in an extended antimicrobial spectrum (1x and 0.1x combined), it represents the most significant growth inhibition caused by the antimicrobial exposure throughout the entire spectrum. Even though there was one categorical susceptibility reporting error in the 1x curve in Fig 5D, all combined categorical susceptibility reports were correct for all conditions. The purpose of the combined GIC reporting from a dual-kinetic-curve response is to only report the maximum growth inhibition and discard GIC reporting errors due to very high or very low microbial loads.

**Table 1.**
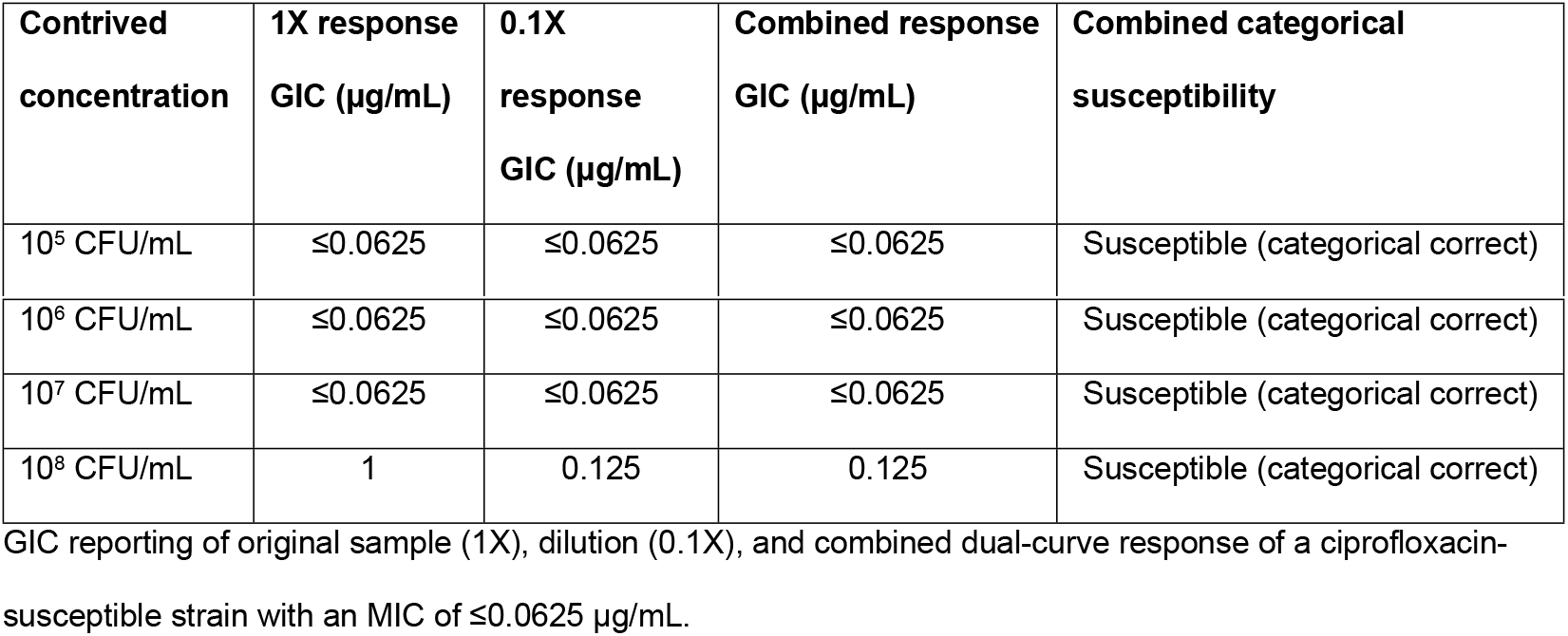
Ciprofloxacin growth inhibition concentration reporting for Fig 5.

| Contrived concentration | 1X response GIC ($\mu\text{g/mL}$) | 0.1X response GIC ($\mu\text{g/mL}$) | Combined response GIC ($\mu\text{g/mL}$) | Combined categorical susceptibility |
| --- | --- | --- | --- | --- |
| $10^5$ CFU/mL | $\leq 0.0625$ | $\leq 0.0625$ | $\leq 0.0625$ | Susceptible (categorical correct) |
| 10<sup>6</sup> CFU/mL | ≤0.0625 | ≤0.0625 | ≤0.0625 | Susceptible (categorical correct) |
| 10<sup>7</sup> CFU/mL | ≤0.0625 | ≤0.0625 | ≤0.0625 | Susceptible (categorical correct) |
| 10<sup>8</sup> CFU/mL | 1 | 0.125 | 0.125 | Susceptible (categorical correct) |

GIC reporting of original sample (1X), dilution (0.1X), and combined dual-curve response of a ciprofloxacin-susceptible strain with an MIC of ≤0.0625 µg/mL.

To evaluate the correlation of the GIC reporting algorithm to the microbial susceptibility and MIC values throughout the physiological range with other antimicrobial classes, the same set of microbial loads in urine (10^5^ to 10^8^ CFU/mL) were repeated with gentamicin in Fig 6 and meropenem in Fig 7. The reported GIC value of each response curve from the dual kinetic (1x and 0.1x) curves from the Maximum Inhibition algorithm is listed in the graph. See S6 and S7 Tables for GIC reporting from all three algorithms. Figs 6 A-D show the transition of GIC reporting for gentamicin from ≤1 μg/mL (susceptible), 4 μg/mL (susceptible), 8 μg/mL (intermediate), to 16 μg/mL (resistant). The categorical susceptibility reporting of “susceptible” was correct over a range of 10^4^ CFU/mL (0.1x of 10^5^ CFU/mL) to 10^6^ CFU/mL (0.1x of 10^7^ CFU/mL). A reporting of 8 μg/mL GIC from 10^7^ CFU/mL (1x of 10^7^ CFU/mL and 0.1x of 10^8^ CFU/mL) is one dilution higher than the MIC of 4 μg/mL, and it is acceptable for essential agreement but a minor error for categorical agreement. The GIC reporting of 16 μg/mL from 10^8^ CFU/mL was a major error. The GC signal levels as shown in S6 Table were saturated at 10,000 nA for 10^7^ and 10^8^ CFU/mL, so the reported GIC value is expected to be higher than the MIC values due to the inoculum effect.

**Fig 6.**
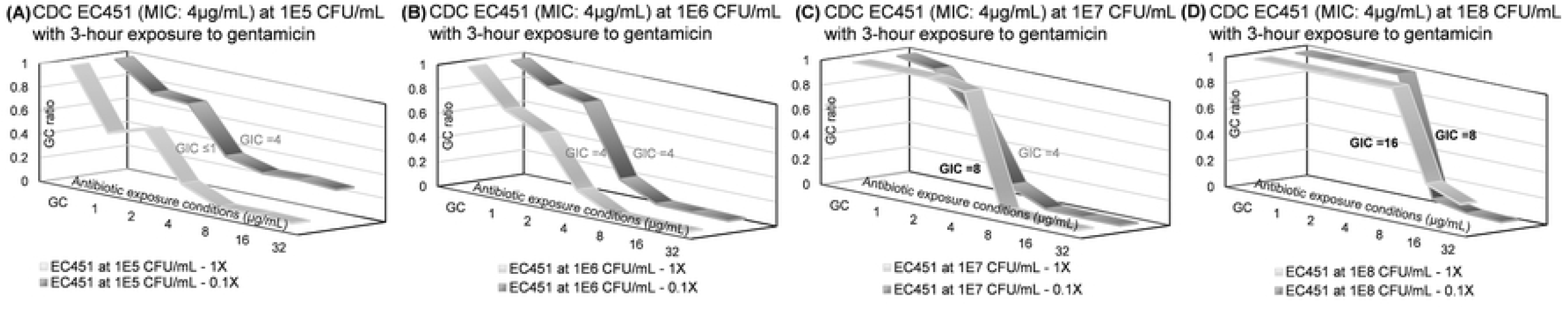
Direct-from-urine gentamicin antimicrobial efficacy profiling with dual kinetic curves on different contrived urine concentrations. Dual kinetic curves for *E. coli* CDC 451 with MIC of 4 μg/mL at starting sample concentrations of (A) 10^5^ CFU/mL, (B) 10^6^ CFU/mL, (C) 10^7^ CFU/mL, (D) 10^8^ CFU/mL. Bolded GIC values indicate incorrect categorical susceptibility.

**Fig 7.**
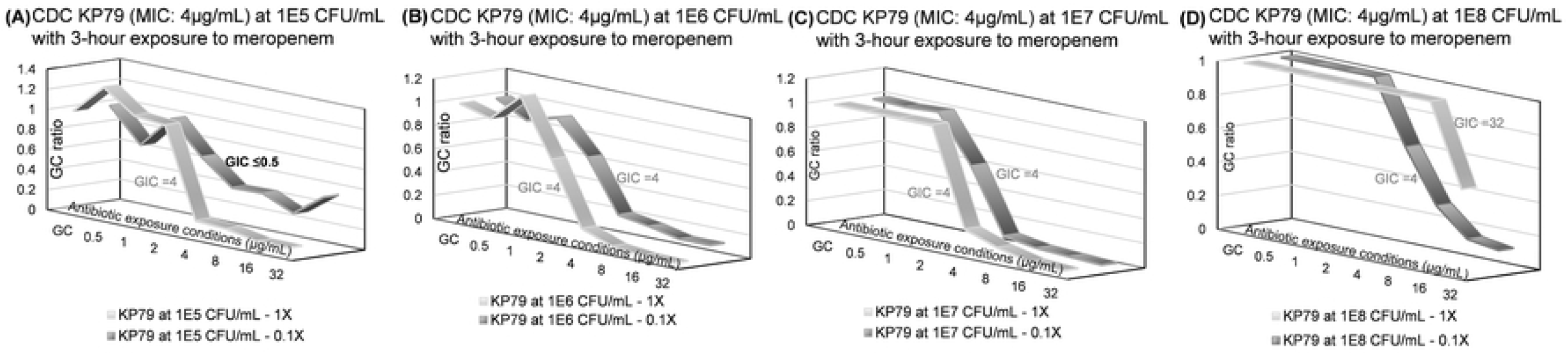
Direct-from-urine meropenem antimicrobial efficacy profiling dual kinetic curves for different starting sample concentrations. Dual kinetic curves for *K. pneumoniae* CDC 79 with MIC of 4 μg/mL at starting sample concentrations of (A) 10^5^ CFU/mL, (B) 10^6^ CFU/mL, (C) 10^7^ CFU/mL, (D) 10^8^ CFU/mL. Bolded GIC values indicate incorrect categorical susceptibility.

Table 2 is the summary of the individual and combined GIC reporting from all contrived concentrations in Fig 6. The MIC value of *E. coli* CDC 451 is 8 μg/mL and therefore intermediate, but our microdilution indicated the MIC was 4 μg/mL and would be classified categorically as susceptible. The combined response GIC for Fig 6C and 6D would be 8 μg/mL to report the maximum growth inhibition, but the combined GIC was adjusted due to signal level saturated at the growth control (GC) and low antibiotic concentrations (1 and 2 μg/mL in 1x response curve in Fig 6C, 1 – 8 μg/mL in 1x response curve and 1-4 μg/mL in 0.1x response curve in Fig 6D). The electrochemical current reading is set to saturate at 10,000 nA to maximize the resolution at lower current readings around the limit of detection. So, the reading would be saturated if the microbial load were too high (>10^8^ CFU/mL). The reported GIC was adjusted one dilution down for every antibiotic concentration reported saturated at 10,000 nA. So the GIC reporting of combined responses from Figs 6C and 6D was adjusted from 8 μg/mL to 4 μg/mL. There were three categorical susceptibility reporting errors in Figs 6C and 6D compared to the microdilution, and all combined categorical susceptibility reporting were correct for all conditions.

**Table 2.** Gentamicin growth inhibition concentration reporting for Fig 6.

| Contrived concentration | 1X response GIC (µg/mL) | 0.1X response GIC (µg/mL) | Combined response GIC (µg/mL) | Combined categorical susceptibility |
| --- | --- | --- | --- | --- |
| 10<sup>5</sup> CFU/mL | ≤1 | 4 | 4 | Susceptible (categorical correct) |
| 10<sup>6</sup> CFU/mL | 4 | 4 | 4 | Susceptible (categorical correct) |
| 10<sup>7</sup> CFU/mL | 8 → 4 | 4 | 4 | Susceptible (categorical correct) |
| 10<sup>8</sup> CFU/mL | 16 → 8 | 8 → 4 | 4 | Susceptible (categorical correct) |

GIC reporting of original sample (1X), dilution (0.1X), and combined dual-curve response of a gentamicin susceptible strain with an MIC of 4 µg/mL.

Similar results were observed for the same study with meropenem in Fig 7. The reported GIC transitioned from ≤0.5 μg/mL (susceptible), 4 μg/mL (susceptible), to 32 μg/mL (resistant). The categorical susceptibility reporting of “resistant” was correct over a range of10^5^ CFU/mL to 10^8^ CFU/mL. A reporting of ≤0.5μg/mL GIC from 10^4^ CFU/mL (0.1x of 10^5^ CFU/mL) was a very major error for categorical agreement, but the GC signal level as shown in S7 Table was 39 nA, which is considered “no growth.” No GIC value would be reported in the case of GC failure (<50 nA).

Table 3 is the summary of the individual and combined GIC reporting in Fig 7. The MIC value of *K. pneumoniae* CDC 79 is 8 μg/mL as listed in CDC AR bank database, but our microdilution indicated the MIC was 4 μg/mL. The combined response GIC for Figs 7A and 7D would be 0.5 and 32 μg/mL, respectively, to report the maximum growth inhibition, but the combined GIC was adjusted due to growth control failure (39 nA for 0.1x of 10^5^ CFU/mL) and the signal level saturated at the growth control (GC) and five low antibiotic concentrations (0.5 to 16 μg/mL in 1x response curve in Fig 7D). Originally, there was only one categorical susceptibility reporting error in Fig 7A, but it would not be reported due to GC fail. All combined categorical susceptibility reports were correct for all conditions.

**Table 3.** Meropenem growth inhibition concentration reporting for Fig 7.

| Contrived concentration | 1X response GIC (µg/mL) | 0.1X response GIC (µg/mL) | Combined response GIC (µg/mL) | Combined categorical susceptibility |
| --- | --- | --- | --- | --- |
| 10<sup>5</sup> CFU/mL | 4 | ≤0.5 | 4 | Resistant (categorical correct) |
| 10<sup>6</sup> CFU/mL | 4 | 4 | 4 | Resistant (categorical correct) |
| 10<sup>7</sup> CFU/mL | 4 | 4 | 4 | Resistant (categorical correct) |
| 10<sup>8</sup> CFU/mL | 32 → 20 | 4 | 20 | Resistant (categorical correct) |

GIC reporting of original sample (1X), dilution (0.1X), and combined dual-curve response of a meropenem-resistant strain with an MIC of 4 µg/mL.

After the initial validation of the presented microbial growth inhibition response curves to antibiotic exposure conditions with CDC clinical strains, we conducted a pilot feasibility study on blinded urine specimens from NYPQ. De-identified clinical remnant specimens were shipped overnight to GeneFluidics for testing as described above, and the summary of combined categorical susceptibility is detailed in Table 4. Sample #7 was positive for *P. aeruginosa* but when tested with the assay produced a GC fail. Subculture of NYPQ sample #7 on Chromagar plate indicated two separate strains, so the original specimen might have been a polymicrobial infection or there was contamination during sample collection or testing. The species-specific susceptibility reporting would require the pathogen identification (ID) sensor chip with complementary oligonucleotide probes against each target pathogen, which is outside the scope of this study. All other nine specimens were reported correctly to match the categorical susceptibility verified by NYPQ. All individual and combined GIC reports are listed in S8 Table. Because NYPQ’s AST panel tests levofloxacin (LEV) instead of ciprofloxacin (CIP) for the class of fluoroquinolones, the GIC reporting of the CIP susceptibility for Samples 1, 4, and 6 were compared to the categorical susceptibility as determined by the reference broth microdilution method for ciprofloxacin. Susceptibility data show levofloxacin to be less potent than ciprofloxacin against gram-negative pathogens such as *Pseudomonas aeruginosa* and certain *Enterobacterales* [54–55]. If a pathogen is susceptible to levofloxacin, it might not be susceptible to ciprofloxacin as seen in Sample 4. However, if a pathogen is resistant to ciprofloxacin, it is most likely to be resistant to levofloxacin as seen in Sample 6.

**Table 4.** Summary of direct-from-urine antimicrobial efficacy profiling using de-identified remnant urine specimens from NYPQ.

| Sample Code | Organism | 1X response GIC (µg/mL) | 0.1X response GIC (µg/mL) | Combined response GIC (µg/mL) | NYPQ reported susceptibility (MIC: µg/mL) | Combined categorical susceptibility |
| --- | --- | --- | --- | --- | --- | --- |
| 0001 | *Citrobacter koseri* | 0.125 | <0.06 | <0.06 | CIP susceptible<br>($\leq 0.0625$) | CIP Susceptible<br>(categorical correct) |
| 0002 | *Escherichia coli* | <0.5 | 0.5 | <0.5 | MEM susceptible<br>($\leq 0.25$) | Susceptible<br>(categorical correct) |
| 0003 | *Enterobacter cloacae* complex | 16 | 16 | 16 | GEN resistant<br>($\geq 16$) | Resistant<br>(categorical correct) |
| 0004 | *Escherichia coli* | 0.5 | 0.25 | 0.5 | CIP susceptible<br>(0.5) | CIP Intermediate<br>(categorical correct) |
| 0005 | *Serratia marcescens* | <0.5 | No growth | <0.5 | MEM susceptible<br>($\leq 0.25$) | Susceptible<br>(categorical correct) |
| 0006 | *Klebsiella pneumoniae* | >4 | >4 | >4 | CIP resistant<br>($> 4$) | CIP Resistant<br>(categorical correct) |
| 0007 | *Pseudomonas aeruginosa* | No growth | No growth | | MEM resistant<br>($\geq 32$) | No growth |
| 0008 | *Proteus penneri* | <1 | <1 | <1 | GEN susceptible<br>($< 1$) | Susceptible<br>(categorical correct) |
| 0009 | *Citrobacter koseri* | <1 | <1 | <1 | GEN susceptible<br>($\leq 1$) | Susceptible<br>(categorical correct) |
| 0010 | *Escherichia coli* | 32 | 32 | 32 | GEN resistant<br>($\geq 16$) | Resistant<br>(categorical correct) |

## Discussion

To date, PCR-based pathogen identification can be performed in less than 30 minutes, but no phenotypic AST exists that can be performed within a reasonable time frame (in hours) directly from clinical samples in clinical microbiology laboratory settings. Schoepp et. al. demonstrated that AST results can be obtained by using benchtop digital nucleic acid LAMP quantification of DNA replication to measure the phenotypic response of fast-growing *E. coli* present within clinical urine samples exposed to an antibiotic for 15 min, but only highly resistant or susceptible strains were selected for testing [56]. For slow-growing pathogens, a longer antibiotic-exposure incubation would be required. Khazaei et. al. demonstrated that quantifying changes in RNA signatures instead of DNA replication resulted in significant shifts (>4-fold change) in transcription levels within 5 min. of antibiotic exposure [57–58]. However, there was a wide range of control:treated ratio (C:T ratio) dispersion from highly susceptible strains with MICs at least seven 2-fold dilutions below the resistant break point. The C:T ratio can change from 2 to 6 with 8 strains with an MIC of 0.015 μg/mL and one strain with an MIC of 0.03 μg/mL, while the C:T ratio separation between the resistant and susceptible populations is only about 0.4. This indicates the limitation in clinical settings when not all susceptible strains have MICs that low.

While the concept of direct-from-specimen AST or antimicrobial efficacy profiling is appealing, there are significant challenges to this approach. The first challenge is that most growth-based susceptibility testing requires a standardized inoculum where a known concentration of organism is used for AST. In routine testing, the organism concentration is fixed, and it may be significantly higher than what is encountered in a clinical specimen which may be used for direct inoculation. An exception may be the urine culture, where patients with real infections commonly have more than 10^5^ CFU/ml. Mezger et al. published a proof-of-concept study in which urine was used as an inoculum for rapid AST [59]. This method employed a brief incubation period (~120 min) followed by quantitative PCR designed to quantify growth. Pilot experiments showed that the assay was able to accurately determine *E. coli* susceptibility to ciprofloxacin and trimethoprim within 3.5 h, however the susceptibility profiling algorithm was not correlated to CLSI M100 categorical reporting. This challenge is addressed by assessing susceptibility response dynamic trends at three different bug/drug ratios by inoculating the raw specimens in two dilutions as detailed above. The second challenge is to provide susceptibility profiling equivalent to AST reported by a clinical microbiology lab with >95% categorical agreement. The third challenge is the need to ensure pathogens are isolated from clinical samples to allow for retesting, confirmation of phenotypic testing (e.g., AST) or epidemiological studies. This challenge will be addressed by setting aside the remainder of specimens for QC or archiving purposes.

Despite being recognized as the standard quantitative index of antimicrobial potency, the MIC is subject to several limitations. It is determined only at the end-point between 16 and 24 hours, at a low initial bacterial inoculum (i.e. 3 to 5 colonies usually in the absence of resistant populations), and utilizes constant (i.e. static) antibiotic concentrations [60]. Therefore, the MIC neither provides information on the time-course of bacterial killing nor on emergence of resistance [61–65]. Several static and dynamic *in vitro* and *in vivo* infection model studies have been demonstrated for analysis and interpretation of *in vitro* efficacy results of antimicrobial drugs as an alternative to MIC reporting [66–71]. These experimental models provide a wealth of time-course data on bacterial growth and killing, but have not adopted into a diagnostic test directly from clinical specimens [72].

An ideal growth inhibition spectrum can fit concentration-responses in sigmoidal curves that are symmetrical about its inflection point and flattened on both ends with statistical fluctuations as shown in Figs 5–7. The left plateau represents insignificant grow inhibition under antibiotic exposures below the MIC, and the right plateau represents significant grow inhibition above the MIC. The inflection point indicates the concentration at which antimicrobial potency lies midway between non-inhibited growth (left plateau) and total inhibited growth (right plateau), and the slope of the tangent to the curve at the inflection point is a measure of the antimicrobial intensity.

With predetermined concentrations of antibiotics in each growth well, the effectiveness of the antibiotics increases and lowers the rate of viability; and this is reflected in the growth control (GC) ratio, which would be negatively correlated with the instantaneous mortality rate. Therefore, the concentration at the inflection point or GIC should increase when the microbial load in the clinical specimen is higher. This agrees with the studies published by other groups [73–76]. Based on this hypothesis, we developed a direct-from-specimen microbial growth inhibition test with two dilutions from unprocessed clinical specimens (1x and 0.1x) as inoculums for two sets of antibiotic exposure stripwells with one GC and seven antibiotic concentrations each to generate a microbial growth inhibition spectrum. As the drug concentration increases, the probability that drug molecules reach a lethal concentration increases as a function modeled by a smooth sigmoidal curve. Since the microbial load in the clinical specimen is unknown, the coverage of this spectrum is designed to capture the inflection point within the whole range of physiological conditions. The GC well of each stripwell serves two purposes: (1) GIC adjustment based on the microbial load under no antibiotics, and (2) quality control to eliminate the data set if there is no growth due to microbial load below limit of detection (LoD). A tentative algorithm has been developed to identify the antibiotic concentration at the inflection point adjusted by the microbial load from the signal level from GC wells, and the reported GICs were compared to MIC from reference methods or FDA cleared systems.

## Supporting information

**S1 Table. Clinical isolate counts with strain # and antimicrobial tested (MIC).** On-scale strains in bold.

**S2A Table. GIC reporting values for Fig 2A.** Ciprofloxacin GIC reporting with three algorithms for *E. coli* CDC 69 with a MIC of ≤ 0.0625 μg/mL and *K. pneumoniae* CDC 79 with an MIC of >8 μg/mL for Fig 2A.

**S2B Table. GIC reporting values for Fig 2B.** Ciprofloxacin GIC reporting with three algorithms for *E. coli* CDC 69 with a MIC of ≤ 0.0625 μg/mL and *K. pneumoniae* CDC 79 with a MIC of >8 μg/mL for Fig 2B.

**S2C Table. GIC reporting values for Fig 2C.** Meropenem GIC reporting with three algorithms for *E. coli* CDC 77 with a MIC of ≤ 0.12 μg/mL and *E. coli* CDC 55 with an MIC of > 8 μg/mL.

**S3A Table. GIC reporting values for Fig 3A.** Gentamicin GIC reporting with three algorithms for *E. coli* CDC 1 with an MIC of 4 μg/mL.

**S3B Table. GIC reporting values for Fig 3B.** Meropenem GIC reporting with three algorithms for *E. coli* CDC 85 with an MIC of 1 μg/mL.

**S3C Table. GIC reporting values for Fig 3C.** Ciprofloxacin GIC reporting with three algorithms for *K. pneumoniae* CDC 80 with an MIC of 0.5 μg/mL.

**S3D Table. GIC reporting values for Fig 3D.** Ciprofloxacin GIC reporting with three algorithms for *K. pneumoniae* CDC 80 with an MIC of 0.5 μg/mL.

**S4A Table. GIC reporting values in three algorithms for Fig 4A.**

**S4B Table. GIC reporting values in three algorithms for Fig 4B.**

**S5 Table. GIC reporting values for Fig 5.** Ciprofloxacin GIC reporting with three algorithms for *E. coli* CDC 69 with a MIC of ≤ 0.0625 μg/mL.

**S6 Table. GIC reporting values for Fig 6.** Gentamicin GIC reporting with three algorithms for *E. coli* CDC 451 with a MIC of 4 μg/mL.

**S7 Table. GIC reporting values for Fig 7.** Meropenem GIC reporting with three algorithms for *K. pneumoniae* CDC 79 with a MIC of 4 μg/mL.

**S8 Table. GIC reporting of NYPQ blinded clinical specimens.**

